# Nanobody immunolabelling and three-dimensional imaging reveals spatially restricted LYVE1 expression by kidney lymphatic vessels in mice

**DOI:** 10.1101/2024.08.09.607309

**Authors:** Eva Maria Funk, Daniyal J Jafree, Nils Rouven Hansmeier, Clàudia Abad Baucells, Rose Yinghan Behncke, Gideon Pomeranz, Maria Kolatsi-Joannou, William J Mason, Dale Moulding, Lauren G Russell, Sascha Ulfers, Laura Wilson, David A Long, René Hägerling

## Abstract

Lymphatic vessels are complex three-dimensional (3D) structures that facilitate tissue fluid clearance and regulate immune responses in health and inflammatory contexts. Recent advances in wholemount immunolabelling and 3D imaging have provided insights into organ-specific heterogeneity of lymphatic structure and function. However, the visualisation of lymphatic vessels deep within an intact organ remains a challenge. We hypothesised that nanobodies, single-domain antibodies raised in camelid species, would result in improved labelling of lymphatics in intact mouse organs, without loss of information due to organ sectioning or inadequate penetration. We generated and characterised nanobody clones targeting lymphatic vessel endothelial hyaluronan receptor 1 (LYVE1), a marker of lymphatic capillaries. Nanobodies were superior at penetrating whole mouse organs and enhanced labelling of lymphatics compared with a conventional anti-LYVE1 polyclonal antibody. Utilising this new tool, we found that kidney lymphatics; an organ in which labelling of lymphatics is challenging, have spatially restricted LYVE1 expression compared with lymphatics of skin, heart, and lung. The timing of this LYVE1 spatial restriction coincides with the early postnatal period in mice. Our findings highlight an unexpected, organ-specific characteristic of kidney lymphatic vessels, whilst providing a novel experimental tool for characterisation, isolation, or perturbation of lymphatic vessels in health and disease.

## INTRODUCTION

The lymphatic vasculature comprises a complex network of blind-ended vessels with roles in tissue fluid homeostasis and the clearance of immune cells and macromolecules to modulate inflammatory responses (Oliver et al. 2020). Lymphatics are present in nearly all adult organs, and have been implicated in a wide range of pathological contexts including cancer metastasis (Le et al. 2016) and anti-tumour immunity (Karakousi, Mudianto, and Lund 2024), cardiovascular diseases (Klaourakis, Vieira, and Riley 2021), and autoimmunity (Bouta et al. 2018). The range of diseases in which lymphatics have been implicated highlight organ-specific heterogeneity in their structure and function (Ulvmar and Mäkinen 2016). Furthermore, evidence for lymphatic heterogeneity comes from organ specific roles including intestinal dietary lipid absorption (Nurmi et al. (2015), cerebrospinal fluid clearance (Ahn et al. 2019) and regulation of intraocular drainage (Park et al. 2014). However, due to their complex, three-dimensional (3D) structure and relative rarity compared to other cell types, traditional histological and imaging techniques fall short in accurately identifying and outlining the structural features of lymphatic vessels. This limitation poses a significant challenge in understanding organ-specific heterogeneity of lymphatic vasculature, underscoring the urgent need for improved technologies for lymphatic labelling and imaging (Liu, Glaser, et al. 2021).

Recently, whole-mount immunofluorescence has enabled the analysis of the complete lymphatic vascular network of mouse organs within which these vessels are superficially located, such as the meninges (Antila et al. 2017; Louveau et al. 2015), skin (Zhang et al. 2022; Karaman et al. 2015), and heart (Trincot et al. 2019; Vieira et al. 2018). In certain organs, such as the kidney (Donnan, Kenig-Kozlovsky, and Quaggin 2021) (Jafree and Long 2020) and liver (Jeong et al. 2023; Jeong, Tanaka, and Iwakiri 2022), lymphatics are located deep within the tissue, necessitating optical clearing techniques that homogenise tissue refractive index (Ueda et al. 2020). This approach has been successfully used to characterise the emergence of lymphatics in mouse embryogenesis (Hägerling et al. 2013) and appreciate structural changes to lymphatics in mouse models of kidney disease (Liu, Hiremath, et al. 2021), human lymphedema (Hägerling et al. 2017) and transplant rejection (Jafree et al. 2022). However, without using alternative strategies such as genetic reporter mice (Redder et al. 2021), lengthy labelling times (Cai et al. 2023) or specialised perfusion equipment (Mai et al. 2023), wholemount immunolabelling is limited by the depth of penetration of conventional IgG antibodies into intact organs.

We hypothesised that harnessing camelid-derived single-domain antibodies, termed nanobodies, with a small size of approximately 15 kDa, would enable deep tissue penetration, and improved epitope detection compared with conventional IgG antibodies (Mitchell and Colwell 2018; Dumoulin et al. 2002; Hassanzadeh-Ghassabeh et al. 2013), making them ideally suited for immunolabelling lymphatics in intact organs. To this end, we developed nanobodies targeting lymphatic vessel endothelial hyaluronan receptor 1 (LYVE1), a CD44-like transmembrane glycoprotein (Johnson et al. 2021) and widely used lymphatic capillary marker (Banerji et al. 1999). We validated these anti-LYVE1 nanobodies before utilising them for 3D imaging and quantitative analysis, finding that nanobody labelling is superior to conventional immunolabelling for visualisation of lymphatic capillaries in intact mouse organs. In so doing, we reveal a novel organ-specific molecular feature of lymphatics, finding that kidney lymphatic capillaries have a spatially restricted expression of LYVE1 compared to these vessels in other organs, a phenomenon which occurs from an early postnatal stage.

## RESULTS

### Generation and production of nanobodies targeting mouse LYVE1

We selected LYVE1 as a target for nanobody generation, primarily as it is a candidate marker for lymphatic capillaries across organs which, if labelled successfully, would enable 3D quantitative analysis of organ-specific lymphatic vascular networks in the body (Jafree et al. 2019). To generate nanobodies (**Fig. 1A-C**), peripheral blood lymphocytes were isolated from llamas that were repeatedly immunised with recombinant mouse LYVE1 protein fused with a polyhistidine (His10) tag (**Fig. 1D**). cDNA from all nanobody sequences present in peripheral blood lymphocytes, regardless of specificity for LYVE1, were amplified by PCR to generate a library of candidate nanobody clones. Nanobodies specific to mouse LYVE1 were enriched by biopanning (**Fig. 1E**). Based on ELISA data, six highly specific anti-LYVE1 nanobodies were chosen for production (**Fig. 1F**). The sequences of these six nanobodies were cloned into an expression vector carrying a 6xHistidine tag and, after successful bacterial production, the nanobody clones were purified by His-affinity purification. We confirmed that successful purification was achieved with a single band observed between 11-17 kDa on Coomassie-blue stained sodium dodecyl sulfate–polyacrylamide gel electrophoresis (SDS-PAGE) gels (**Fig. 1G**) and following western blotting using anti-His Ab (**Fig. 1H**). A summary of our workflow can be found in **Fig. S1**.

**Figure 1:**
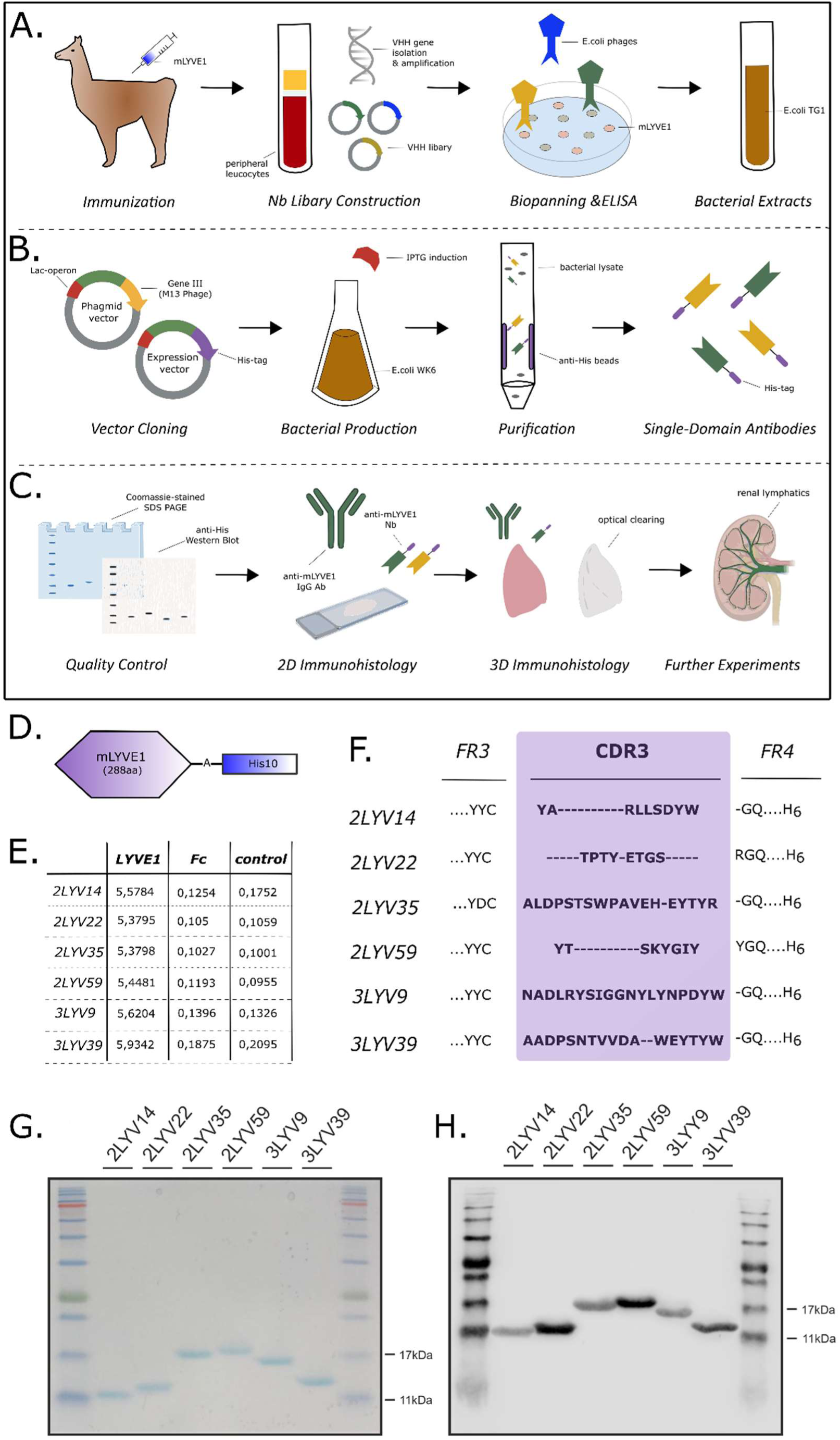
Generation and production of an anti-mouse LYVE1 nanobody. To generate the anti-mouse LYVE1 nanobody (**A**), a recombinant mouse LYVE1 protein, consisting of 288 amino acids (aa), fused to a His10 tag (**D**) was repeatedly injected into Ilama *(Immunisation)*. Peripheral blood lymphocytes were harvested, and total RNA extracted for first-strand cDNA synthesis. Nanobody-encoding sequences were amplified and cloned into phagemid vector pMECs and transformed into E. coli TG1, resulting in a nanobody library of 10^9^ independent transformants *(Library Construction)*. Next, anti-mouse LYVE1-specific nanobodies were enriched by phage display and biopanning on a solid phase coated mouse LYVE1. LYVE1-binding nanobodies were further analysed towards binding performance by whole-cell phage ELISA (**E**). Wells assessed were coated with (1) mouse LYVE1 with His10 tag and mouse LYVE1 fused to human IgG1 Fc, (2) the human IgG1 Fc (Fc) or (3) blocking buffer only (control). The values indicate specificity of clones, which was used to determine which to produce. *(Biopanning & ELISA)*. Finally, bacteria carrying the anti-mouse LYVE1 nanobody plasmid were used for further downstream steps *(Bacterial Extracts)*. For production (**B**) nanobody sequences were cloned into pHEN6c expression vector carrying an additional 6x His *(Vector Cloning)*. Amino acid sequence of complementary determining region (CDR) 3 with accompanying Framework region (FR) 3 and 4 of six anti-mouse LYVE1 nanobody produced are displayed in (**F**). Next, WK6 E. coli carrying pHEN6c-nanobody plasmids were cultured and production was induced by addition of isopropyl-ß-D-1-thiogalactopyranoside (IPTG) *(Bacterial Production)*. Bacterial cells were then lysed and nanobodies separated from other cell components by His affinity purification *(Purification)*. Finally, His-tagged LYVE1 nanobodies were ready for further experiments. *(Single Domain Antibodies)*. To validate (**C**) LYVE1 nanobodies, sodium dodecyl sulphate-polyacrylamide gel electrophoresis (SDS-PAGE) with Coomassie blue staining and anti-His western blotting were performed. Both confirmed single protein bands of the expected nanobody size of 11-17kDa (**G**). Anti-His western blotting found His-positive proteins (**H**). Taken together, these results acted as a confirmation of successful nanobody production and purification *(Quality Control)*. To assess the suitability of nanobodies for immunohistology, nanobodies were primarily tested in two-dimensional (2D) histological sections using anti-mouse LYVE1 IgG antibody as a control (**Fig. S2**) *(2D Immunohistology)*. Next, nanobodies were validated for use in wholemount immunolabelling and optical clearing (Fig. 2) *(3D Immunohistology).* Subsequently, nanobodies were utilised to investigate the presence of LYVE1 in various conditions *(Further Experiments)*.

### Validation of anti-mouse LYVE1 nanobodies for 2D and 3D imaging

Next, we evaluated the performance and specificity of the six anti-mouse LYVE1 nanobody clones. Initially, we performed conventional two-dimensional (2D) immunofluorescence, by labelling cryosections of C57BL/6 mice at embryonic day (E) 14.5 with either anti-LYVE1 nanobodies or a commercially available anti-LYVE1 IgG antibody and comparing patterns of expression. All six nanobody clones exhibited staining patterns overlapping with that of the LYVE1 antibody, successfully capturing primitive network of jugular lymphatic sacs that give rise to systemic lymphatic vasculature (Yang and Oliver 2014) (**Fig. S2A**). To assess the specificity of anti-LYVE1 nanobodies in a 3D context, we performed nanobody labelling of optically-cleared lungs from reporter mice (n=3) carrying Cre and an eGFP cassette within the endogenous *Lyve1* locus (*Lyve1^eGFP-Cre^*) (Pham et al. 2010). *Lyve1^eGFP-Cre^* mice at postnatal day (P) day 28 exhibited GFP expression in lymphatic endothelial cell nuclei, which overlapped with the cell surface labelling of our anti-LYVE1 nanobodies (**Fig. 2A**). These findings validate that the newly produced nanobodies recapitulate endogenous LYVE1 expression and demonstrate their specificity and suitability for 3D imaging of optically-cleared biological tissue.

**Figure 2:**
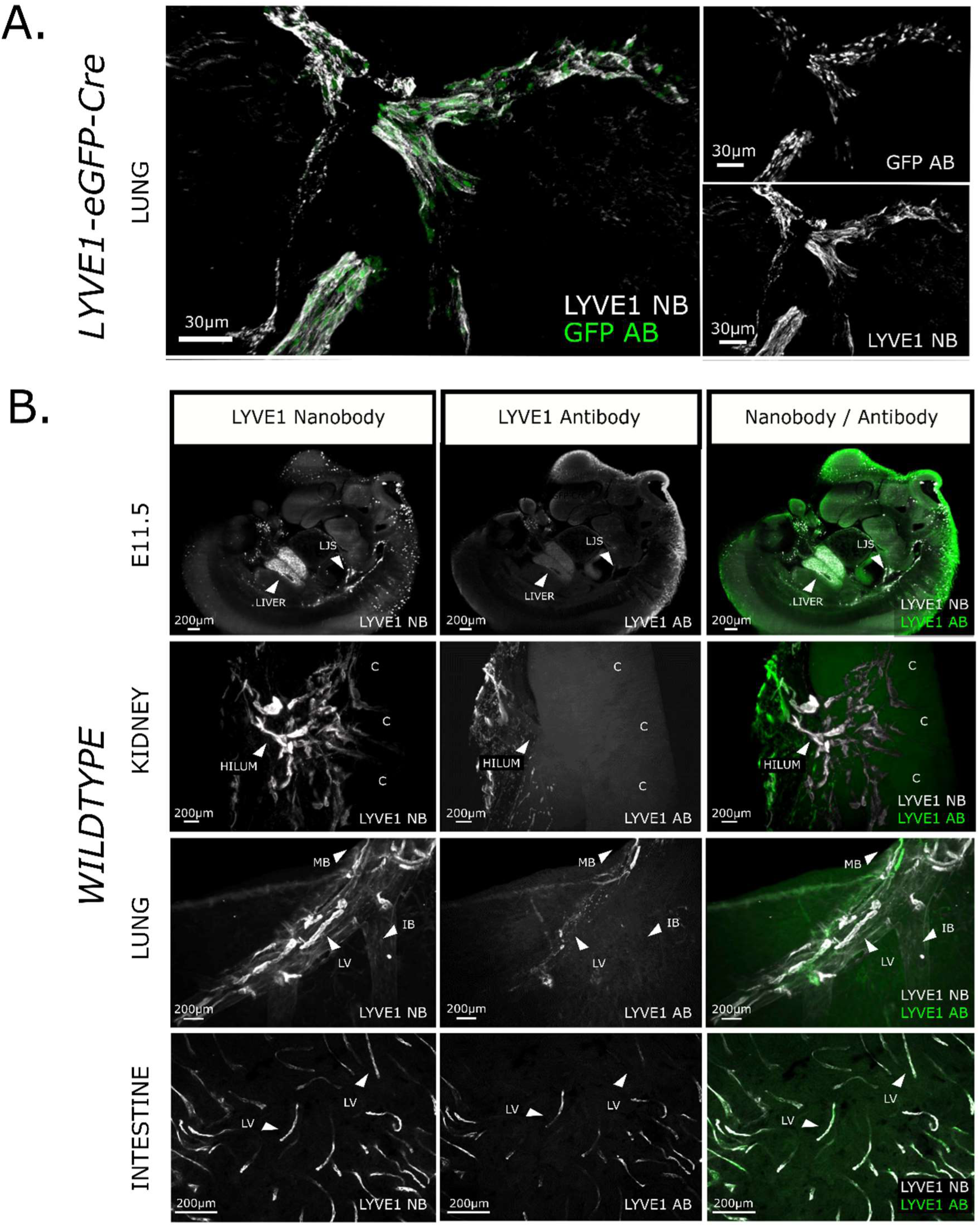
Characterisation and evaluation of an anti-mouse LYVE1 nanobody for 3D imaging of intact mouse organs. (**A**) Co-staining of anti-mouse LYVE1 nanobodies (white) and anti-GFP IgG antibodies (green) in P28 *Lyve1^eGFP-Cre^* mouse lung tissue (n=3) visualised by confocal microscopy. Both anti-GFP IgG antibodies and LYVE1 nanobodies visualise a large lymphatic vessel. While the anti-GFP IgG antibody shows the expected nuclear staining pattern, the LYVE1 nanobodies visualise lymphatic endothelial cell surfaces. This not only reinforces the specificity of anti-mouse LYVE1 nanobodies shown previously in 2D histology (**Fig. S2**), but also further supports the feasibility of using LYVE1 nanobodies in tissue cleared specimens. The scale bar represents 30 µm. (**B**) Evaluation of the efficacy of LYVE1 nanobodies in wholemount immunostaining of entire tissue samples. This panel compares the efficacy of LYVE1 nanobodies (white) to LYVE1 IgG antibodies (green) controls using light sheet microscopy in various intact mouse organs. In E11.5 mouse embryos (n=6), the jugular lymphatic sac (JLC) is distinctly visible in the LYVE1 nanobody channel, while the antibody channel exhibits a weak signal with a halo around the embryo surface, indicating limited IgG antibody penetration. Furthermore, the LYVE1+ liver sinusoidal endothelial cells are visualised in great detail by the LYVE1 nanobodies, which the LYVE1 IgG antibodies lack. In P28 kidney (n=6), nanobodies efficiently label lymphatic vessels beyond the superficial hilum vessels and further into the kidney, whereas IgG antibodies mainly visualises lymphatic vessels on the hilum surface. The nanobodies allow evaluation of interlobular lymphatic vessels extending deep into the cortex (C). P28 lung lobes (n=6) exhibit a higher number of deeper lymphatic vessels (LV) and increased bronchial autofluorescence in the nanobody channel. While IgG antibodies primarily visualised vascular structures overlying the main bronchus (MB) near the tissue surface, nanobodies additionally reveal vascular structures accompanying the intermediate bronchus (IB). In the small intestine (n=6), confocal visualisation shows increased staining of lymphatic vessels (LV) in the nanobody channel. In comparison, IgG antibodies detect less LV, at lower intensity or only partially. The scale bar represents 200 µm.

### Improved detection of organ lymphatic vessels using anti-mouse LYVE1 nanobodies

Next, we tested whether using nanobodies targeting LYVE1 provides improvements in the detection of lymphatics in intact, optically-cleared mouse organs, compared with conventional IgG antibodies. To do this, we performed wholemount immunolabelling of wildtype mouse embryos (n=6) and a range of visceral organs at P28 (n=6 per organ). We harvested intestine, a hollow viscous tissue amenable to wholemount techniques (Zarkada et al. 2023), alongside lung (Stump et al. 2017) and kidney (Jafree and Long 2020) where lymphatics are located deep within the tissue, necessitating optical clearing prior to 3D imaging. Tissues were labelled using three anti-mouse LYVE1 nanobody clones (2LYV14, 2LYV22, 2LYV35) which exhibited the highest signal-to-background ratio in 2D staining experiments (**Fig. S2B**), or a commercially available anti-LYVE1 IgG antibody.

In all tissues examined, the commercial anti-LYVE1 IgG antibodies exhibited superficial staining, with weak immunoreactivity of lymphatic vessels deep within each tissue. E11.5 embryos additionally exhibited a halo of non-specific staining around the perimeter of the tissue. By contrast, the nanobodies penetrated deeper in both the embryos and organs, enhancing the detection of lymphatics in the E11.5 embryo and intact P28 intestine, lung, and kidney (**Fig. 2B**). Within E11.5 embryos, the nanobodies enabled visualisation of known LYVE1^+^ structures such as the jugular lymphatic sac or LYVE1^+^ liver sinusoidal endothelial cells (Guo et al. 2022). Within the intact P28 kidney, nanobody labelling revealed interlobular lymphatic vessels extending from the hilum to the organ’s cortex (Donnan, Kenig-Kozlovsky, and Quaggin 2021; Jafree and Long 2020), whereas the conventional anti-LYVE1 IgG antibodies only labelled superficial hilar vessels. Within the P28 lung parenchyma, lymphatic vessels running alongside the bronchi could be visualised in greater detail with nanobody labelling as compared with conventional antibodies. Intestinal lymphatics also exhibited more widespread labelling with the nanobody approach (**Fig. 2B**).

To determine if non-lymphatic LYVE1^+^ structures were being captured, we further immunolabelled E18.5 kidneys with the myeloid marker, F4/80. LYVE1^+^ macrophages have been reported in embryonic kidneys (Lee et al. 2011) and indeed, using nanobody labelling, F4/80^+^ LYVE1^+^ cells were detected within E18.5 kidney (**Fig. S3A**). In these experiments, we also demonstrated a further methodological advantage over conventional IgG antibodies in that the incubation period with anti-LYVE1 nanobodies could be reduced to as little as four hours to achieve effective staining, whereas the equivalent staining with conventional anti-LYVE1 IgG antibodies was only visible after 48 hours (**Fig. S3B**). Overall, these findings show that anti-LYVE1 nanobodies are more efficacious than IgG antibodies for wholemount immunofluorescence, optical clearing and 3D imaging of lymphatics within mouse embryos and adult organs.

### Quantitative comparison reveals spatially restricted expression of LYVE1 as an organ-specific feature of kidney lymphatics

Having successfully developed anti-LYVE1 nanobodies for wholemount immunofluorescence in mouse tissues, we hypothesised this novel reagent would enable us to examine LYVE1 expression and lymphatic architecture across a range of postnatal mouse organs. Therefore, we assessed heart, lung, skin and kidney at P28 in C57BL/6 wildtype mice (*n*=8 mice per group), a timepoint where we reasoned that the developmental remodelling of lymphatics would be complete (Klaourakis, Vieira, and Riley 2021). Focussing on imaging the parenchyma of each tissue to capture lymphatic capillaries, we undertook comparative 3D image analysis by utilising anti-LYVE1 nanobodies and an anti-podoplanin (PDPN) IgG antibody. PDPN is expected to label all postnatal lymphatic endothelial cells (Breiteneder-Geleff et al. 1997; Guo et al. 2022; Dick et al. 2022), serving as means of comparing the number of lymphatic capillaries which also expressed LYVE1. Being a conventional IgG antibody, PDPN antibodies did not fully penetrate all organs, thus LYVE1 and PDPN vessel structures were extracted separately, binarized and analysed for geometric properties including vessel branch volume, length or mean radius using open-source 3D analysis software (Bumgarner and Nelson 2022). In regions of tissues fully penetrated by both immunolabels, PDPN should capture all lymphatic vessels, thus LYVE1^+^ vessels were examined relative to their PDPN^+^ counterparts (**Fig. S4**).

We inspected 3D images from P28 heart, lung, and skin, finding consistent overlap between PDPN and LYVE1 expression profiles in all organs examined (**Fig. 3A-C**). Accordingly, no significant difference between the relative volume of LYVE1^+^ vessels and PDPN^+^ vessels were found in the skin (p=0.15), heart (p=0.15) or lung (p=0.88) (**Fig. 3E-G**). However, the kidney showed a clear discrepancy. Analysing over 30 imaging volumes of individual vessel segments within the kidney’s parenchyma, we consistently identified kidney lymphatic vessels expressing PDPN, but lacking expression of LYVE1 (**Fig. 3D**). Quantitative analysis substantiated our findings, revealing a statistically significant difference (p=0.03) between the relative volume of LYVE1^+^ vessels and that of PDPN^+^ vessels (**Fig. 3H**).

**Figure 3:**
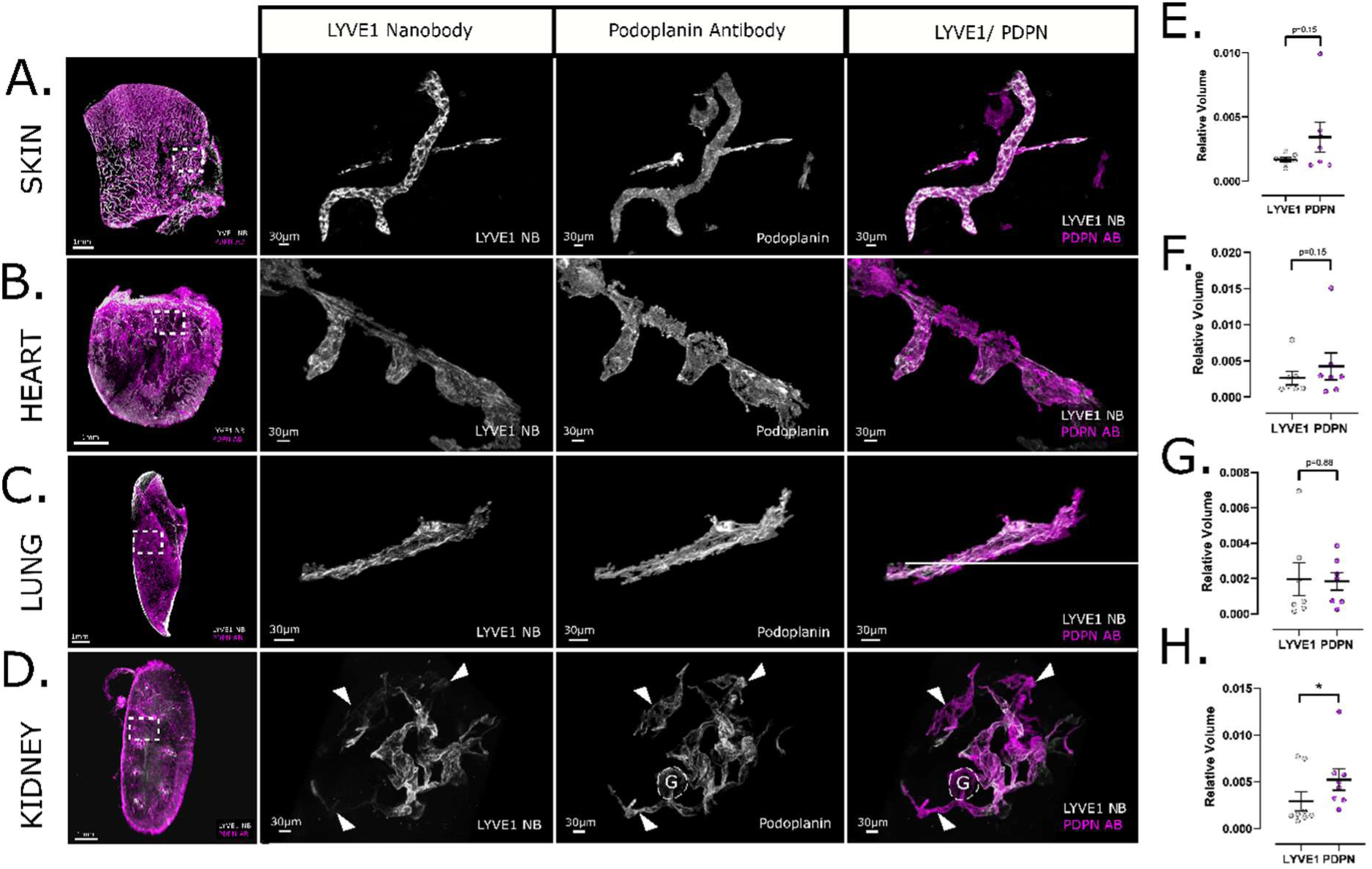
3D imaging of nanobody-stained mouse organs reveals spatially restricted expression of LYVE1 as an organ-specific of kidney lymphatic vasculature. Co-staining of anti-mouse LYVE1 nanobodies (white) and anti-PDPN IgG antibodies (magenta) in C57BL/6 wild-type P28 organs by confocal microscopy. Each channel is displayed separately in grey scale. The scale bar in the overview is 1 mm and 30 µm for regions of interest. (**A-D**) Comparison of the vascular structures visualised by anti-mouse LYVE1 nanobodies and anti-PDPN IgG antibodies in P28 skin, heart, and lung (**A-C**) reveal full coherence between the lymphatic vessel markers LYVE1 and PDPN, as expected. To distinguish lymphatic vessels from bronchi, bronchial structures are outlined with dotted lines (**B**). In renal tissue (**D**) an incoherence with a lack or reduced levels of LYVE1 in individual vessel segments was found (Fig. 2A-C). Arrows indicate LYVE1^−^ vessel segments. PDPN is a marker for both lymphatic endothelial cells and podocytes, which form the glomerulus. Consequently, the PDPN channel also visualises glomeruli (G), as indicated by the dotted lines. (**E-H**) Quantitative analysis of LYVE1^+^ and PDPN^+^ lymphatic vessels in P28 organs. Each data point represents the average relative volume of 3-4 regions of interest imaged per animal. The error bar shows the standard error of means. An analysis of single regions of interest per data point can be found in the supplements (**Fig. S6A-D**) Vessel volume has been adjusted to the overall sample volume. In P28 skin (n=6), a paired student’s t-test found no significant difference (p=0.15) between the relative volumes of PDPN and LYVE1 (mean difference = 0.0017, t=1.626, df=6) (**E**). Similarly, in P28 heart (n=7), no significant difference was observed (p=0.15, mean difference = 0.0016, t=1.662, df=6) (**F**). In P28 lung (n=7), the difference was also not significant (p=0.88, mean difference = −0.00012, t=0.1459, df=6) (**G**). However, in P28 kidneys (n=8), a significant difference was found between the relative volumes of PDPN and LYVE1 (p=0.039, mean difference = 0.002311, t=2.528, df=7) (**H**).

The expression of LYVE1 in adjacent vessel branches suggests that the lack of expression of LYVE1 by kidney lymphatics is not an artefact of limited immunolabel penetration into the tissue. These LYVE1^−^ PDPN^+^ vessels were always continuous with LYVE1^+^ PDPN^+^ lymphatics, demonstrating that they were part of the lymphatic network and not anatomically separate. Moreover, the arrangement of LYVE1^−^ PDPN^+^ vessels was non-hierarchical, with lack of LYVE1 expression by lymphatics captured both in the hilum and cortex of the kidney, suggesting that this feature does not originate from a pre-collecting or collecting vessel phenotype (Ulvmar and Mäkinen 2016; Mäkinen et al. 2005). Thus, a combination of nanobody labelling, 3D visualisation and quantitative analysis demonstrated an organ-specific spatial restricted expression of LYVE1 by kidney lymphatics.

### Temporal decline in LYVE1 expression by kidney lymphatics occurs during postnatal maturation

Finally, given the organ-specific nature of restricted LYVE1 expression by lymphatics in the kidney, we sought to gain insight into the dynamics of LYVE1 expression over the course of kidney development and maturation. We characterised and compared 3D imaging of wildtype kidney lymphatics at five timepoints: E18.5, P1, P5, P28 and P90. E18.5 (n=8) was chosen as an embryonic timepoint at which we have previously characterised the spatial relationships of kidney lymphatics, and during which LYVE1 is expressed by all bona fide lymphatic vessels (Jafree et al. 2019). Between P1 (n=7) and P5 (n=8), the development of nephrons; the functional units of the kidney, has been reported to reach cessation (Li et al. 2021) with P28 (n=8) representing early adulthood and P90 (n=6) representing a mature stage by which time mouse kidneys have reached their full size.

At E18.5, all PDPN^+^ lymphatic vessels were found to express LYVE1 and, accordingly, quantitative analysis demonstrated no significant differences between LYVE1^+^ vessel and PDPN^+^ vessel volume (p=0.96) (**Fig. 4A-B**). Likewise, there was no significant difference in LYVE1^+^ vessel and PDPN^+^ vessel volume at P1 (p=0.33) (**Fig. 4A,C**), or P5 (p=0.15), albeit there were some visible regions containing LYVE1^−^ PDPN^+^ lymphatic vessels at this early postnatal timepoint (**Fig. 4A,D**). Thereafter, there was a striking decrease in LYVE1^+^ regions of PDPN^+^ lymphatic vessels as the mice aged. (**Fig. 4A**). By P28, there was a significant difference between LYVE1^+^ vessel and PDPN^+^ vessel volume (p=0.03) (**Fig. 3H**), consistent with findings at P90 (p=0.04) (**Fig. 4E**). Non-parametric multiple comparisons were applied to evaluate LYVE1^−^ vessels across the five different timepoints. Using this approach, no significant differences could be found between time points E18.5 and P1 (p=0.95), nor between P28 and P90 (p=0.77) or P5 and P28 (p=0.68) (**Fig. 4F**). However, in an analysis of individual regions of interests across all animals, a significant difference between P5 and P28 was detected (p= 0.03) (**Fig. S5G**). The most pronounced difference in this analysis within maturation stages occurred between P1 and P5 (p=0.002) (**Fig. 4F**). In summary, our 3D quantitative analysis of LYVE1 expression during kidney lymphatic vessel remodelling revealed that the organ-specific feature of restricted LYVE1 expression manifests during the organ’s early postnatal maturation, occurring most rapidly between P1-P5.

**Figure 4:**
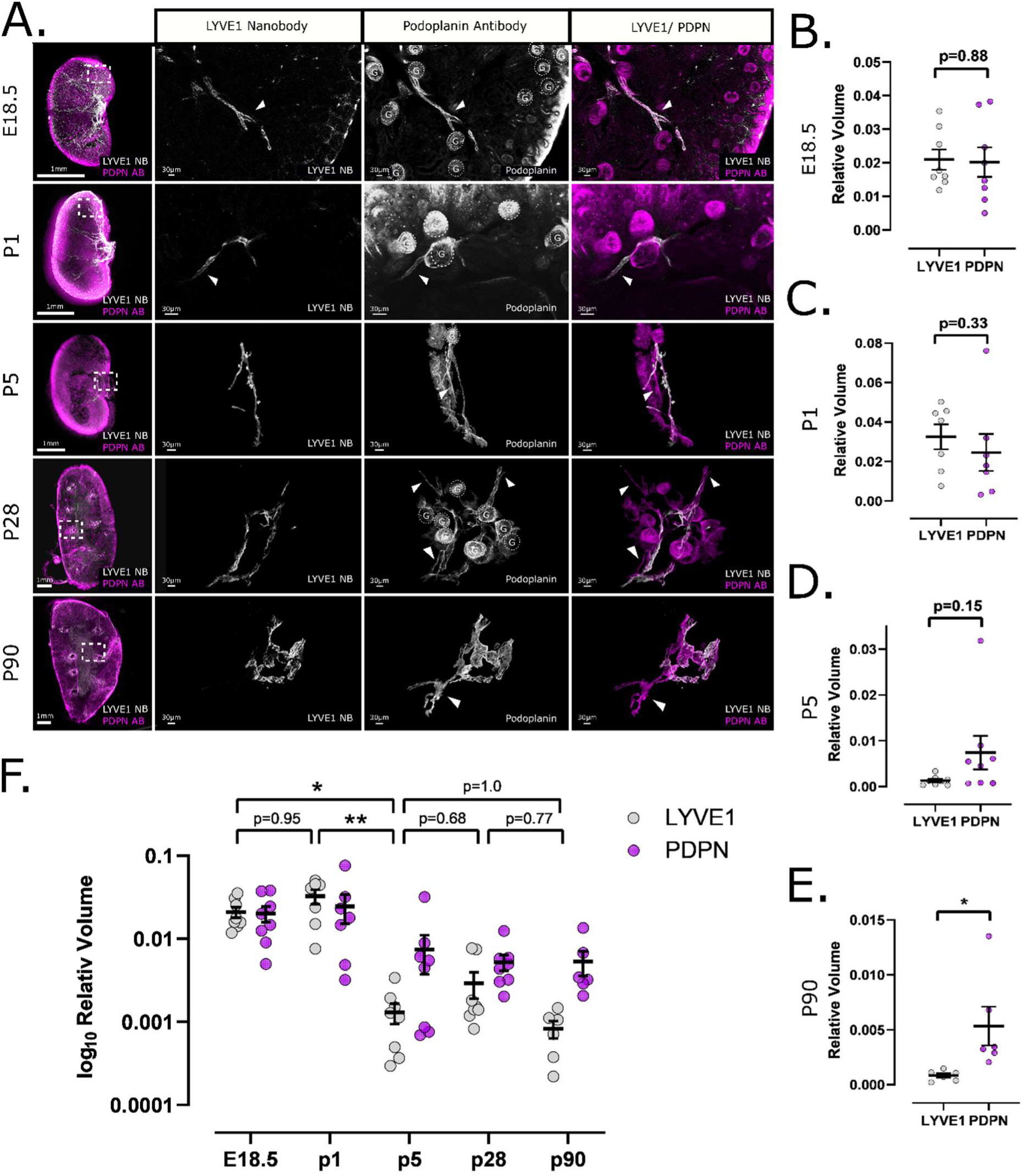
Spatial restricted expression of LYVE1 expression by kidney lymphatics occurs within the postnatal period. (**A**) Anti-mouse LYVE1 nanobodies (white) and anti-PDPN IgG antibodies (magenta) staining of C57BL/6 wild-type kidney tissue sections at various time points visualised by confocal microscopy. Each channel is displayed separately in grey scale. Regions of interest were visualised in areas of the kidney tissue sample as indicated in the overview image. At E18.5 and P1, both LYVE1 and PDPN equally visualise vascular structures in the sample and no incoherence can be detected. However, different staining patterns between LYVE1 and PDPN were observed in P5 kidneys, although incoherence occurred in a smaller number of vessels and regions of interest compared to P28 kidney tissue (Fig. 3D). P90 renal lymphatics showed a similar number of LYVE1^−^ areas, particularly in vessels of decreasing size. Arrows indicate LYVE1^−^ vessel segments. Glomeruli (G) visualised by the PDPN IgG antibody are outlined by dotted lines in the grey scale image of PDPN. The scale bar is 1 mm, 30 µm respectively. (**B-F**) Quantitative analysis of LYVE1^+^ and PDPN^+^ lymphatics vessel in kidneys. Each datapoint represents the average relative volume of 3-4 regions of interest imaged per kidney, except for E18.5 and P1 as kidneys were imaged as a complete unit. Error bar shows standard error of means. Vessel volume has been adjusted to the overall sample volume. An analysis of single regions of interest per data point can be found in the supplements (**Fig. S6E-F**). In E18.5 kidneys (n=8), a paired student’s t-test found no significant difference (p=0.88) between the relative volumes of PDPN and LYVE1 (mean difference = −0.0007419, t=0.1533, df=7) (**B**). Similarly, in P1 kidneys (n=7), no significant difference was observed (p=0.33, mean difference = −0.007961, t=1.046, df=6) (**C**). In P5 kidneys (n=8), the difference was also not significant (p=0.15, mean difference = 0.006101, t=1.642, df=7) (**D**). However, in P90 kidneys (n=6), a significant difference was found between the relative volumes of PDPN and LYVE1 (p=0.049, mean difference = 0.004502, t=2.574, df=5) (**E**). (**F**) Statistical analysis of LYVE1 volume dynamics throughout kidney maturation was conducted using nonparametric multiple comparisons for relative contrast effect testing (estimation method = global pseudo ranks, type of contrast = Tukey, confidence level = 95%) (Konietschke et al. 2015) across five developmental timepoints, revealing a significant overall alteration (p=0.0004, quantile=2.74). No significant differences were observed between E18.5 and P1 (p=0.95, estimator=0.603, lower=0.292, upper=0.848, statistic=0.879), P5 and P28 (p=0.68, estimator=0.344, lower=0.131, upper=0.646, statistic=-1.423), P5 and P90 (p=1.0, estimator=0.50, lower=0.221, upper=0.779, statistic=0.00), or P28 and P90 (p=0.77, estimator=0.344, lower=0.117, upper=0.674, statistic=-1.295). However, a significant decline was observed between P1 and P5 (p=0.002, estimator=0.076, lower=0.013, upper=0.345, statistic=-3.698). Additionally, significant differences were noted between non-chronological timepoints, such as E18.5 and P28 (p=0.0004, estimator=0.027, lower=0.002, upper=0.241, statistic=-4.045), P1 and P28 (p=0.003, estimator=0.071, lower=0.011, upper=0.356, statistic=-3.575), and P1 and P90 (p=0.0013, estimator=0.042, lower=0.004, upper=0.296, statistic=-3.792). Statistical analysis of single regions of interest can be found in **Fig. S6G.**

## DISCUSSION

In this study, we successfully generated and validated novel anti-LYVE1 nanobodies to improve 3D imaging of lymphatic capillaries in mice. Our approach enables faster and more effective wholemount immunostaining of intact murine organs, facilitating both qualitative and quantitative analysis of organ-specific lymphatic architecture. Testament to the potential of this novel tool for discovery, we found that LYVE1 expression was spatially restricted in lymphatic vessels of the kidney as compared to other organs, and that this phenomenon manifests during the organ’s postnatal maturation, coinciding with the cessation of organogenesis in the kidney. In developing a reagent for 3D lymphatic imaging, we have therefore highlighted an organ-specific feature of lymphatics in the kidney and demonstrate the importance of exercising caution when using single markers, such as LYVE1, to discriminate lymphatics in uncharacterised tissues.

Previous reports have harnessed the beneficial properties of single-domain antibodies for 3D imaging, specifically their small size, improved epitope detection compared with conventional IgG antibodies (Muyldermans 2013), high solubility, heat stability, chemical resistance (Dumoulin et al. 2002) and cost-effectiveness (Muyldermans 2021). Recent examples of 3D imaging utilising nanobodies include labelling human dermal vasculature (Hansmeier et al. 2022), capturing GFP^+^ cells in whole mice (Cai et al. 2023) or characterising the murine placental vasculature (Freise et al. 2023). Such nanobodies surmount the challenges of using conventional IgG antibodies for immunolabelling, including long incubation times or specialised equipment being required for tissue penetration (Mai et al. 2023). To our knowledge, this represents the first nanobody that binds to lymphatics. LYVE1 was selected given its reported widespread expression on lymphatic capillaries (Oliver et al. 2020), and its amenability to successful nanobody production given its simple structure and limited number of post-translational modifications compared to other lymphatic markers such as PDPN. The nanobody clones presented in this report have other applications beyond 3D imaging, including flow cytometry and cell isolation, live imaging or functional blocking experiments in mice (Babamohamadi et al. 2024; Pymm et al. 2021).

Having generated and validated anti-LYVE1 nanobodies, we demonstrate whole organ imaging of lymphatic vascular networks. Although previous reports have utilised LYVE1 as a marker in isolation to capture lymphatic vessels (Baranwal et al. 2021; Lee et al. 2011), nanobody labelling revealed that LYVE1 expression is spatially restricted by lymphatics of the kidney from an early postnatal stage. Given the location of these LYVE1^−^ PDPN^+^ capillaries and their absence in other organs, we provide evidence for an organ-specific feature of lymphatics in the mouse kidney. We have recently shown that these findings also translate to human tissues, as 3D imaging of human kidneys demonstrates regions of lymphatic vessels with LYVE1^−^ PDPN^+^ capillaries (Jafree et al. 2022). Furthermore, transcriptional comparison using single-cell RNA sequencing data highlights that approximately 30% of lymphatic cells in the kidney lack *LYVE1* expression (Jafree et al. 2022). This demonstration of intra-organ molecular heterogeneity of lymphatics may also apply to other contexts (Ulvmar and Mäkinen 2016), such as the nasal lymphatics, within which a population of atypical LYVE1^−^ lymphatic capillaries was also identified (Hong et al. 2023).

Our study is not without limitations. Firstly, our validation of nanobodies in 3D imaging data is performed with co-labelling of conventional IgG antibodies. As a result, experiments are limited by the penetration depth of full-size IgG antibodies, necessitating separate analysis of PDPN^+^ vessel and LYVE1^+^ vessel volumes for quantitative analysis. Secondly, the nanobodies generated bind to mouse LYVE1, but to our knowledge, do not exhibit reactivity with human LYVE1. Moreover, our study does not address the functional significance of lymphatic heterogeneity, which is out of the scope of this technical report. CD44^+^ immune cells enter lymphatics *via* hyaluronan-mediated binding to LYVE1 on lymphatic endothelium (Johnson et al. 2017; Johnson et al. 2021), thus the limited expression of LYVE1 on kidney lymphatics may have implications for renal immune cell trafficking (Kitching and Hickey 2022; Riedel, Turner, and Panzer 2021). In the mouse nasal mucosa, single-cell transcriptomics indicates that LYVE1^−^ lymphatic cells are enriched for expression of molecules involved in stimulation or regulation of immune responses (Hong et al. 2023), suggesting that this subpopulation may be directly involved in local nasal immunity. Finally, deciphering the origins of lymphatic heterogeneity remains a subject of ongoing debate. Our findings support a developmental origin of restricted LYVE1 expression, as we observe the greatest increase in the proportion of LYVE1^−^ vessel volume during a postnatal window during which kidney organogenesis ceases. Thus, differences in organ-specific lymphatic development, including paracrine or physical signals from tissue progenitors, transcriptional enhancers or repressors and alternative cellular lineages are all potential contributors to structural, molecular and functional heterogeneity of lymphatic vessels (Jafree et al. 2021).

In conclusion, we report the generation of the first nanobodies targeting lymphatic vessels. We find that anti-LYVE1 nanobodies represent a promising and simple-to-use tool to structurally profile organ lymphatics in mouse, rendering the complex technique of 3D imaging more widely accessible within the rapidly evolving field of lymphatic biology. Testament to the utility of these nanobodies, we add to the evidence that lymphatic vessels within certain organs possess unique anatomical and molecular properties, with an atypical LYVE1^−^ lymphatic capillary profile in the mouse kidney. Overall, we anticipate nanobodies will contribute to the suite of novel experimental tools advancing the understanding of lymphatic heterogeneity in health and disease.

## MATERIALS AND METHODS

### Llama immunization and nanobody library construction

For nanobody generation, a recombinant LYVE1 protein, consisting of 288 amino acids, fused to a His10 tag was expressed in human embryonic kidney 293 cells. The recombinant protein was injected at days 0, 7, 14, 21, 28, and 35 into a Ilama (*Lama glama*). After immunization, 100 ml of anticoagulated blood were collected from the Ilama on day 40, 5 days after the final antigen injection. Peripheral blood lymphocytes were isolated and total RNA extracted for cDNA first-strand synthesis using oligo(dT) primers. Sequences encoding variable nanobodies were subsequently amplified by PCR, digested with the restriction enzyme *SapI*, and finally cloned into the *SapI* site of the phagemid vector pMECs. Electrocompetent E. coli TG1 (60502, Lucigen, Middleton, WI, USA) were then transformed with VHH sequence harbouring pMECS vectors, resulting in a nanobody library comprising 10^9^ independent transformants. This process has previously been described in detail by (Vincke et al. 2012).

### Biopanning and identification of anti-mouse nanobodies specific to LYVE1

Phage enrichment and biopanning were carried out as detailed by (Vincke et al. 2012). Briefly, the previously constructed nanobody library was panned for 3 rounds on solid phase coated with mouse LYVE1 (100 µg/ml in 100 mM NaHCO_3_ pH 8.2) yielding a 400-fold enrichment of antigen-specific phages after the 3^rd^ round of panning. A total of 380 colonies were randomly selected and assayed for mouse LYVE1-specific antigens by ELISA, again using mouse LYVE1 and additionally mouse LYVE-1 fused to human IgG1 Fc at the C-terminus (50065-M02H, Sino Biological, Beijing, China). To exclude potential nanobody clones binding to human IgG1 Fc, human IgG1 Fc (10702-HNAH, Sino Biological) was utilized as a control as well as blocking buffer only (100 mM NaHCO3, with pH 8.2). After comparing binding specificity of different clones with the control values, 278 colonies were identified as positive for mouse LYVE1 binding. Using the sequence data, the number of possible nanobody candidates was further narrowed down to 98, of which 96 were able to specifically bind both mouse LYVE1-His10 and mouse LYVE1 fused to human IgG1 Fc. The remaining unique clones derived from 21 different B cell linages according to their complementary determining region (CDR) 3 groups. Considering the different B cell linages and robustness of ELISA screening data, 6 clones were selected for further experiments.

### Cloning of nanobody sequences in expression vector

To generate 6xHis-tagged nanobodies, nanobody sequences were cloned from the pMECS phagemid vector into the pHEN6c expression vector. Initially, nanobody sequences were amplified by PCR using the following primers:

1. 5’ GAT GTG CAG CTG CAG GAG TCT GGR GGA GG 3’
2. 5’ CTA GTG CGG CCG CTG AGG AGA CGG TGA CCT GGG T 3’

PCR products were purified (QIAquick PCR Purification Kit, 28104, Qiagen, Hilden, Germany) and digested for 20 minutes at 37 °C with *PstI-HF* (R3140, New England Biolabs, Ipswich, MA, USA) and *BstEII-HF* (R3162, New England Biolabs) restriction enzymes, while empty pHEN6c plasmids were concurrently digested with the same restriction enzymes. The empty pHEN6c plasmids, however, were supplemented by 5 units of heat-inactivated (5 minutes at 80 °C) FastAP™ alkaline phosphatase (EF0651, Thermo Fisher Scientific, Waltham, MA, USA). Digestion products were purified (28104, QIAquick PCR Purification Kit, Qiagen) and subjected to T4 DNA ligase-mediated ligation reactions. The ligation reaction was performed at 16 °C for 16 hours using 2.5 units of T4 DNA ligase (M0202, New England Biolabs). Subsequently, newly generated nanobody sequence harboring pHEN6c plasmids were transformed into WK6 E. coli cells (C303006, Thermo Fisher Scientific) and analysed towards correct nanobody sequence integration by Sanger DNA sequencing. To this end, the primers used were as follows:

1. 5’ TCA CAC AGG AAA CAG CTA TGA C 3’
2. 5’ CGC CAG GGT TTT CCC AGT CAC GAC 3’

### Production and purification of anti-LYVE1 nanobodies

WK6 E. coli carrying pHEN6c-Nanobody plasmid were cultivated at 37 °C, shaking in 1L‘Terrific Broth’ medium (2.3 g/L KH2PO4, 16.4 g/L K2HPO4-3H2O, 12 g/L tryptone, 24 g/L yeast extract, 0.4% (v/v) glycerol) complemented with 100 µg/mL ampicillin, 2 mM MgCl2, and 0.1% (w/v) glucose. Nanobody expression was induced at an OD600 of 0.6-0.9 by adding 1nM isopropyl ß-D-1-thiogalactopyranoside. After an incubation period of 16 hours, the nanobodies were extracted by centrifugation (8000× g, 8 min, RT). 18 ml of TES/4 buffer (0.05 M Tris [pH 8.0], 0.125 mM EDTA, 0.125 M sucrose) were added for 1 hour of shaking on ice. The cell suspension was thereafter centrifuged (8000× g, 30 min, 4 °C), and periplasmic protein-containing supernatant was collected. For 6xHis-tagged nanobody extraction, HIS-Select® nickel affinity gel (P6611, Sigma-Aldrich, Darmstadt, Germany) was applied according to the manufacturer’s instructions. The solution was loaded on a PD-10 column (17-0435-01, GE healthcare, Chicago, IL, USA) and nanobodies were eluded via 3 × 1 mL 0.5 M imidazole in phosphate-buffered saline (PBS) (I2399, Sigma-Aldrich). An overnight dialysis (3 kDa MWCO, 66382, Thermo Fisher Scientific) against PBS was carried out to remove undesirable imidazole from the nanobody solution.

### Coomassie-blue stained SDS PAGE and western blotting

To verify nanobody production and pureness, sample protein was separated by molecular weight using established sodium dodecyl sulphate-polyacrylamide gel electrophoresis. For each clone, 5 ug of denatured protein with 0.04% (w/v) OrangeG in ddH2O) were loaded onto the gel alongside a pre-stained protein ladder (ab116029, Abcam, Cambridge, UK). For protein visualisation, gels were treated with Coomassie staining solution (0.1% (w/v), Coomassie Brilliant Blue R-250 (1610400, Bio-Rad Laboratories Inc., Hercules, CA, USA), 50% (v/v) methanol, and 10% (v/v) glacial acetic acid in ddH2O) for 1 hour, followed by incubation with Coomassie detaining solution (50% ddH2O, 40% methanol, 10% acetic acid (v/v/v)). Alternatively, gels were blotted onto a nitrocellulose membrane (1620112, Bio-Rad Laboratories Inc.) and nanobodies were identified using a primary anti-His antibody (12698, Cell Signaling Technology, Danvers, MA, USA) and a secondary anti-rabbit antibody (926-32211, LI-COR Biosciences, Lincoln, NE, USA). Subsequently, blots were analysed by an Odyssey® Fc Imaging System (LI-COR Biosciences).

### Mouse husbandry and acquisition of mouse tissues

C57BL/6 wildtype mice or *Lyve1^Cre-eGFP^* mice (Pham et al. 2010) were maintained in compliance with the UK Animals (Scientific Procedures) Act 1986 and experiments were carried out under a UK Home Office project license (PPL: PP1776587). Further, laboratory mouse work was approved by German federal authorities (LaGeSo Berlin) under the licence number ZH120. Nutrition and water were available to animals *ad libitum*. Adult mice or pregnant mice were sacrificed using CO_2_ inhalation and cervical translocation as a Schedule 1 procedure. The desired organs were obtained from embryonic, juvenile, or adult mice, washed in PBS, and subsequently fixed in 4% (w/v) paraformaldehyde (PFA) in PBS for 4 hours at 4 °C to preserve tissue integrity. After fixation, samples were thoroughly washed in three changes of PBS and stored at 4 °C in PBS containing 0.02% (w/v) sodium azide until further processing.

### Antibodies used for immunofluorescence

The following commercially available antibodies were used: rabbit monoclonal anti-His antibody (12698, Cell Signaling Technologies) [1:200], donkey polyclonal anti-rabbit IgG Alexa Fluor™ 647 antibody (A31573, Invitrogen, Waltham, MA, USA) [1:1000], donkey polyclonal anti-rabbit IgG Highly-Cross-Absorbed Alexa Fluor™ 647 antibody (A32795, Invitrogen) [1:1000], goat polyclonal anti-mLYVE1 (AF2125, R&D Systems, Minneapolis, MN, USA) [1:100], donkey polyclonal anti-goat IgG Alexa Fluor™ 568 antibody (A11057, Invitrogen) [1:1000], donkey polyclonal anti-goat IgG Highly cross-absorbed Alexa Fluor™ 488+ antibody (A32814, Invitrogen) [1:1000], hamster monoclonal anti-Podoplanin (14-5381-82, Invitrogen) [1:200], goat polyclonal anti-Syrian hamster IgG Cross-Absorbed Alexa Fluor™ 546 antibody (A-21111, Invitrogen) [1:1000], chicken polyclonal anti-GFP (ab13970, Abcam) [1:200], donkey anti-chicken Highly cross-absorbed Alexa Fluor™ 488+ antibody (A32931TR, Invitrogen) [1:1000], rat monoclonal anti-mF4/80 antibody (MCA497G, BioRad) [1:50], donkey anti-rat Highly cross-absorbed Alexa Fluor™ 488+ antibody (A48269, Invitrogen) [1:1000]. Nanobodies were detected by anti-His staining in combination with an Alexa Fluor™ dye-conjugated secondary antibody.

### Immunofluorescence staining of cryosections

Snap-frozen 5 μm sections of E14.5 wildtype mice were stained with nanobodies at different concentrations (0.1 μg/ml, 1 μg/ml, 10 μg/ml) and control antibodies as previously described in detail (Hansmeier et al. 2022). Visualisation of representative regions was accomplished using an Axioscope5 fluorescence microscope (Zeiss, Oberkochen, Germany) equipped with a Plan-NEOFLUAR 40x/0.75 objective (Zeiss).

### Wholemount immunofluorescence staining and optical clearing

Mouse organs were either cut into smaller sections measuring 0.5-2 mm or processed as whole specimens. Tissues were dehydrated using an ascending methanol series and bleached overnight at 4 °C in a solution containing 5% H2O2 (VWR Chemicals, Radnor, PE, USA) in methanol. Wholemount staining, embedding and optical clearing was performed as previously described (Hägerling et al. 2013). Intestine samples were not optically-cleared, but mounted in fluorescent medium (S3023, Agilent Technologies, Santa Clara, CA, USA). Nanobody (10 μg/ml), primary IgG Antibodies, and secondary IgG were incubated for various periods of time based on sample size and staining reagent (Nanobodies 2-4 hours, primary antibodies 3-14 days, secondary antibodies 1-7 days).

### Wholemount immunofluorescence staining and optical clearing of E18.5 and p1 specimens

Embryonic and early postnatal kidneys were initially dehydrated and bleached as described above. Following rehydration, samples were permeabilized overnight in a 5% solution of 3-((3-cholamidopropyl) dimethylammonio)-1-propanesulfonate in ddH2O and blocked in PBS supplemented with 0.2% (v/v) Triton X100, 10% (v/v) DMSO, and 6% (v/v) goat serum. Nanobodies (10 μg/ml) and antibodies were diluted in antibody solution (PBS + 0.2% v/v Tween20 + 0.1% v/v heparin solution + 5% v/v DMSO + 3% v/v goat serum + 0.1% w/v saponin) and incubated between 4 hours (nanobodies) and 24 hours (IgG antibodies) at 4 °C. Between staining steps, samples were washed in PBS-Tween20. Clearing was performed as previously described for embryonic kidneys (Jafree et al. 2020)

### Confocal and light sheet imaging

Specimens were imaged using two different microscopy systems. For confocal imaging, an LSM 880 Upright Confocal Multiphoton microscope (Zeiss) equipped with a 20x/NA 1.0 W-plan Apochromat water immersion objective was utilized. A comprehensive description of the setup specific to BABB-cleared specimens for this microscope can be found in previously published work (Jafree et al. 2020). Intact mouse organs were imaged by LaVision Ultramicroscope II with a LaVison BioTec MVPLAPO 2x OC OBE objective. Various magnifications were employed, and image acquisition utilized a step size of 2 μm.

### Image processing and 3D rendering

2D immunofluorescence images were subjected to post-processing using ZEN 3.4 (blue edition) software from Zeiss. Single channels were extracted and saved in TIFF format.

Z-stack datasets were processed and 3D-rendered using Imaris 9.8, a software package provided by Oxford Instruments (Abingdon, UK). To reduce non-specific background signals, single channels were often subjected to the Imaris Surface function. Images of the 3D-rendered data were captured using the Imaris snapshot function and saved in TIFF format.

### Quantification of signal-to-background ratio in 2D immunofluorescence

Signal-to-background ratio was quantified using ImageJ 2.24/1.54f (https://github.com/imagej/ImageJ, (Schindelin et al. 2012)). Ten intensity values were randomly selected in vessel areas and areas that showed no specific staining. The mean values for signal and background areas were calculated and the signal-to-background ratio was determined using the formula: Signal-to-background ratio = mean signal/mean background.

### Quantitative analysis of 3D imaging volumes

The quantitative analysis of three-dimensional volumes commenced with the binarization of single channels using Imaris software (v9.8, Oxford Instruments). The binarization process was conducted using the isosurface rendering function, and for E18.5 and P1 samples, 3D cropping was applied to exclude regions with high PDPN intensity at the kidney surface, thus enhancing binarization accuracy. During surface rendering, the threshold was set to absolute intensity, with manual adjustments to ensure the inclusion of all relevant structures. To eliminate smaller non-specific signals, structures were filtered based on the number of voxels. Following surface generation, PDPN channel-derived non-vascular structures were manually removed using the selection function. Subsequently, the channel of interest was masked using specific settings: constant inside/outside, setting voxels outside the surface to 0.00, and inside the surface to the maximum intensity of the prepared channel. All channels except the newly created masked channel were then deleted, and the single channel was saved in TIFF format. With the binarized files prepared, TIFF files were imported into the VesselVio application (Bumgarner and Nelson 2022). The analysis settings were configured as follows: unit µm, resolution type anisotropic with individual sizes of samples, analysis dimensions 3D, image resolution 1.0 µm^3^, and filters applied to isolate segments shorter than 10.0 µm and purge end-point segments shorter than 10.0 µm. Analysis results were automatically saved in Microsoft Excel files by VesselVio. Among the parameters offered by VesselVio, vessel volume was selected as the parameter for further analysis, as it takes into account both vessel length and width.

### Sample size estimation and statistical analysis

Sample size was estimated based on prior publications of renal developmental studies of the lymphatic vessels (Jafree et al. 2019) and studies investigating the lymphatics in adult mice organs (Vieira et al. 2018; Park et al. 2014) Therefore, we anticipated n=6-8 animals per experiment would be sufficient to power statistical analyses. Statistical analysis was conducted using Prism (v8, GraphPad by Dotmatics, Boston, MA, USA) and RStudio version 12.0 (Posit, Boston, MA, USA). To determine the significance of differences in volume between LYVE1 and PDPN at individual time points and individual organs, a paired student’s *t*-test was carried out. To assess statistical significance across different time points a rank-based approach using the R package nparcomp (Konietschke et al. 2015) with the function mctp1 (multiple comparisons for relative contrast effect testing) was chosen. This approach allows for nonparametric multiple comparisons to evaluate relative contrast effects. A p value of less than 0.05 was considered statistically significant for all tests. Quantitative data was visualised using Prism. All graphical representations present individual data points either by region of interest or by animal, along with the mean and standard error of the mean.

### Preparation of figures and videos

The figures presented in this paper have been prepared using the free software tool Inkscape 1.1.0 (Inkscape Project, 2020) for graphic design and layout. Any modifications made to the images were applied consistently to maintain uniformity across all coherent single and merged channel images.

## Data Availability

The data that support the findings of this study are available from the corresponding author upon reasonable request.

## Authors information

EMF, NRH, DJ, RH and DAL conceived the study. Cloning and production of nanobodies was performed by EMF, NRH and CAB. Acquisition of mouse material, mouse husbandry and immunostaining or histology were performed by EMF, NRH, RYB, ACB, GP, MKJ, WJM, LGR, LW, and SU. Confocal or lightsheet imaging and 3D analysis were performed by EMF, DJ, DM and RYH. Project oversight and supervision was provided by NRH, DJ, RH and DAL. EMF wrote the first draft of the paper, refined by DJ, RH and DAL, and subsequently all authors were involved in revision and preparation of the final manuscript for submission.

## Acknowledgement

We express our gratitude to Gholamreza Hassanzadeh, VIB Nanobody Core, Vrije Universiteit Brussel, Brussel, Belgium, for supervision of llama immunizations and technical expertise. We acknowledge Dr Louise Johnson (University of Oxford, UK) for her technical insights into LYVE1 purification, Dr Robert Lees (UKRI Central Laser Facility, UK) for his lightsheet fluorescence microscopy expertise, Dr. Felix Heymann (Universitätsmedizin Berlin, Germany) for access to confocal microscopy and Prof. Dr. Ansgar Peterson (Julius Wolff Institute, Charité Universitätsmedizin Berlin, Germany) for access to image analysis infrastructure. We thank Stephen Schüürhuis and the Institute of Biometry and Clinical Epidemiology (Charité Universitätsmedizin Berlin, Germany) for statistical consultation services.

## Ethics declaration

All authors declare no conflicts of interest.

---

**Supplementary Figure 1:**
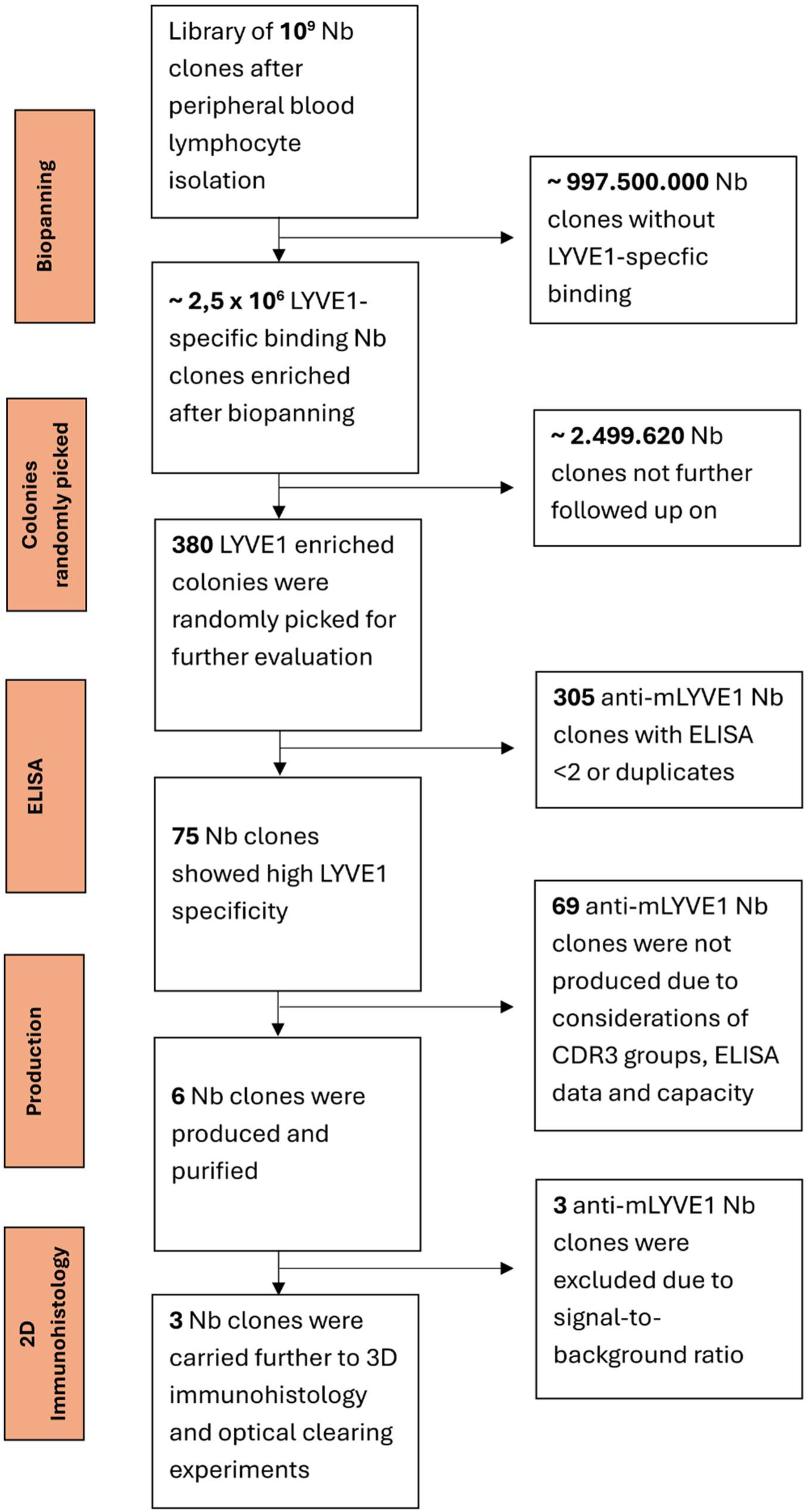
Flowchart of anti-mouse LYVE1 nanobody clone selection process. After initial immunization and establishment of a nanobody library based on peripheral blood leucocytes, a total of 10^9^ potential nanobody clones were available. After 400-fold enrichment by phage display and biopanning of LYVE1-spefic binding clones, 380 clones were randomly selected for ELISA analysis. 305 clones scored <2, which was set as the cut-off for promising mouse LYVE1 high specificity or were duplicates. From the remaining 75 highly LYVE1 specific nanobody clones, due to capacity reasons, merely six clones were selected based on ELISA data and diversity of CDR3 groups for production. All six clones successfully immunolabelled LYVE1 in 2D histological sections, but due to differences in signal-to-background ratio, three clones were finally selected for 3D immunostaining with optical clearing.

**Supplementary Figure 2:**
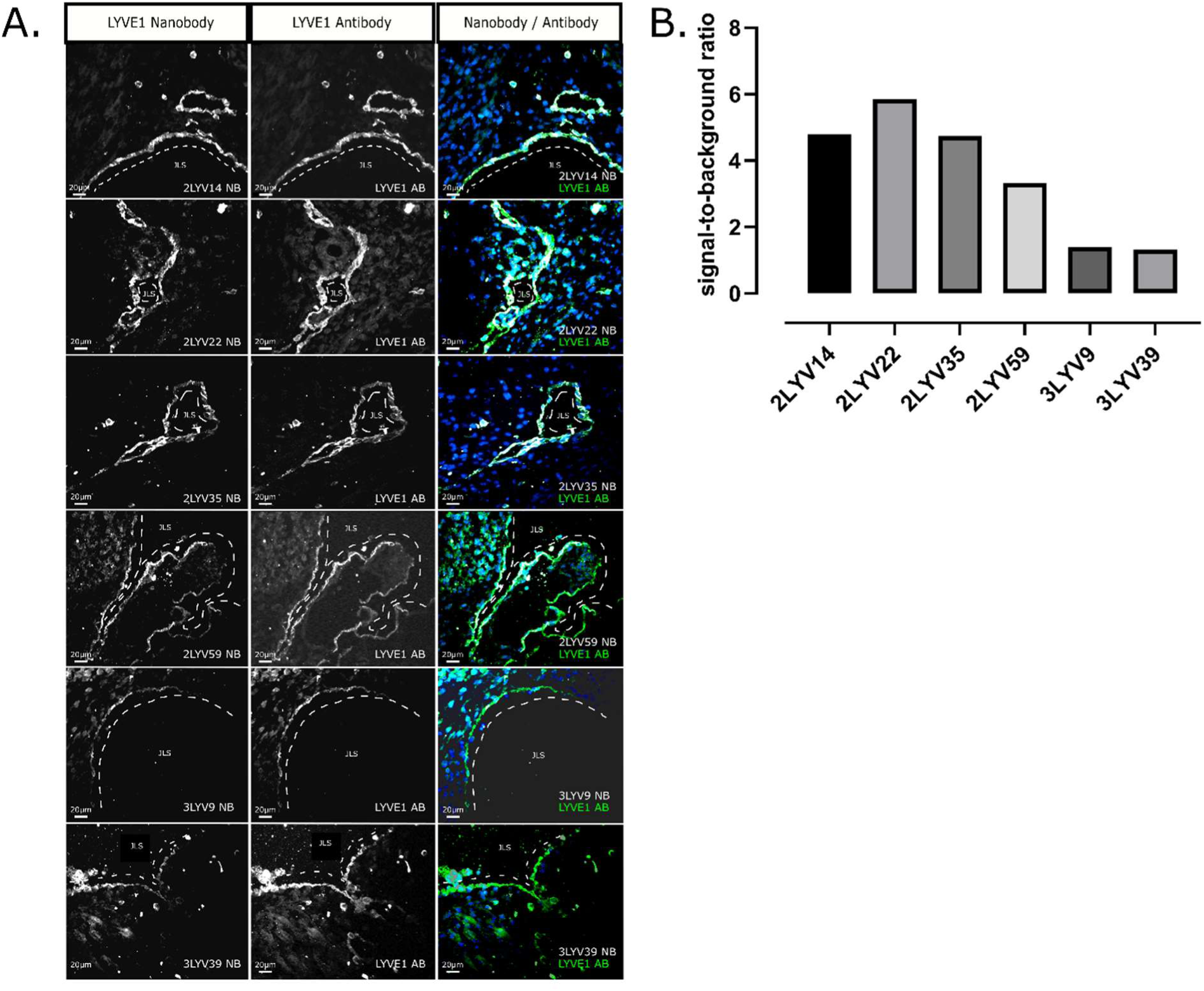
Validation of anti-mouse LYVE1 nanobody for 2D immunolabeling. **(A)** LYVE1 staining of 5 µm cryosections of E14.5 mouse embryos was performed using anti-mouse LYVE1 nanobody clones (white) and IgG antibodies (green), with separate channels displayed in grayscale. The nanobody clones are indicated in each image corner. Hoechst (blue) served as a positive control and is visible in the merged image. Dashed lines outline the jugular lymphatic sac (JLS). Single cells not adjacent to the lymphatic vasculature were detected alongside vascular structures. All six nanobody clones visualised lymphatic structures comparably to IgG controls, often providing improved signal-to-noise ratio. Scale bar 20 µm. **(B)** A quantitative comparison of the signal-to-background ratios for the six anti-mouse LYVE1 nanobody clones in 2D immunofluorescence was performed by selecting ten random values from signal and background areas in ImageJ. The average ratios were 4.8 (2LYV14), 5.9 (2LYV22), 4.7 (2LYV35), 3.3 (2LYV59), 1.4 (3LYV3), and 1.3 (3LYV39). Clones 2LYV14, 2LYV22, and 2LYV35 were selected for 3D immunofluorescence experiments.

**Supplementary Figure 3:**
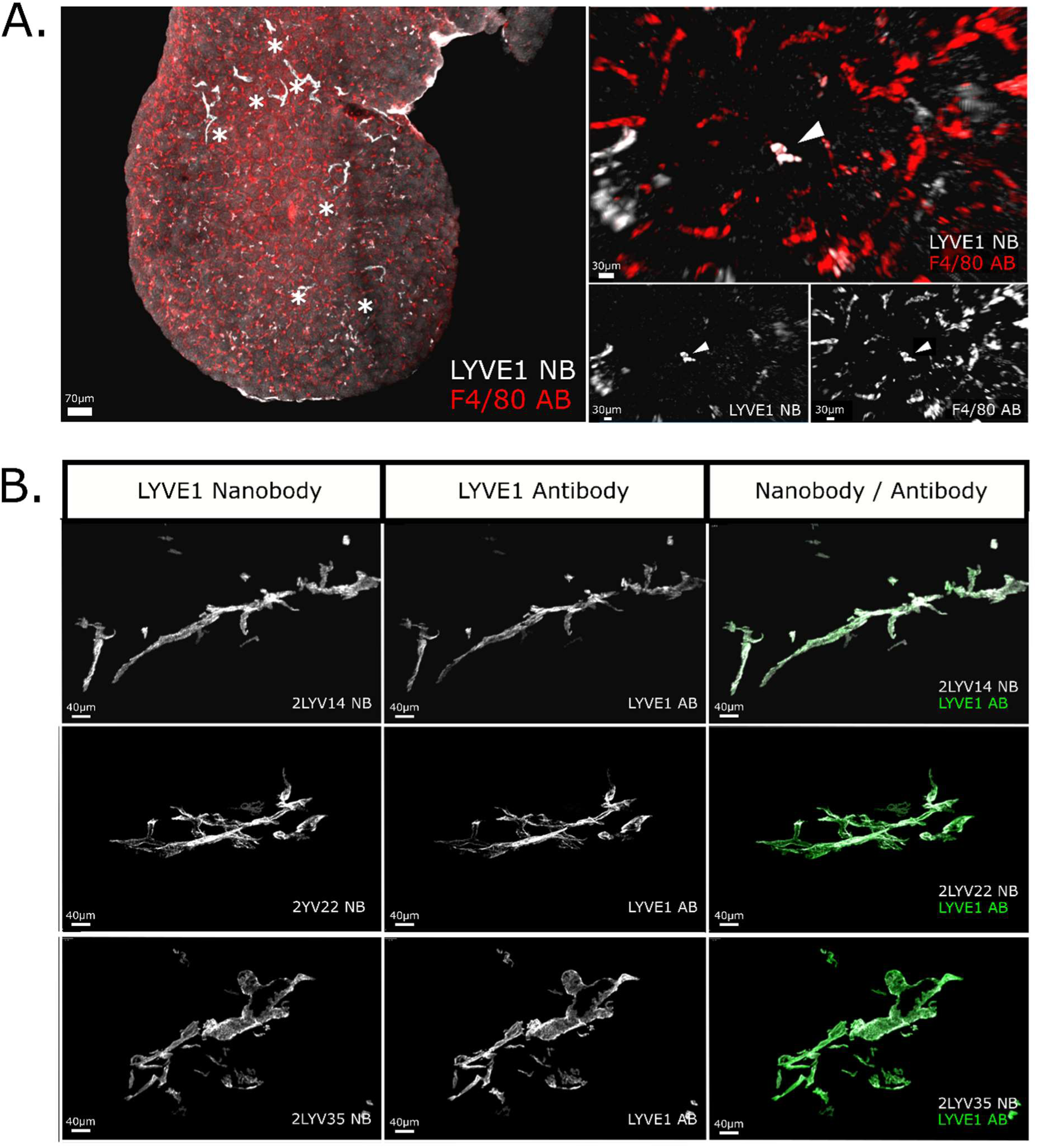
Validation of anti-mouse LYVE1 nanobody in optically-cleared murine kidney after incubation period reduction and validation of myeloid cell expression. **(A)** Co-staining of anti-mouse LYVE1 nanobodies (white) and anti-F4/80 IgG antibodies (red) in C57BL/6 wildtype E18.5 kidney, visualised by confocal microscopy. The macrophage marker F4/80 successfully detected multiple cells within the kidney, while next to lymphatic vessels (*), the anti-mouse LYVE1 nanobodies labelled occasional single cells not connected to the lymphatic vasculature. These cells are co-labelled by anti-F4/80 IgG antibodies, suggesting that the nanobodies are not only capable of labelling LYVE1^+^ lymphatic vessels but can further be used to study LYVE1^+^ macrophage subsets. The scale bar is 70 µm, 30 µm respectively. **(B)** Anti-mouse LYVE1 nanobody clones 2LYVE14, 2LYV22 and 2LYVE35 in wholemount immunostaining of P28 murine kidney sections visualised by confocal microscopy. Control IgG antibodies were incubated for 48 h, in comparison to an incubation period of 4 h for nanobodies. The nanobodies (white) successfully detected the identical biological structures as the IgG antibody control staining (green), despite a noticeable reduction of incubation time by 44h. The scale bar is 40 µm.

**Supplementary Figure 4:**
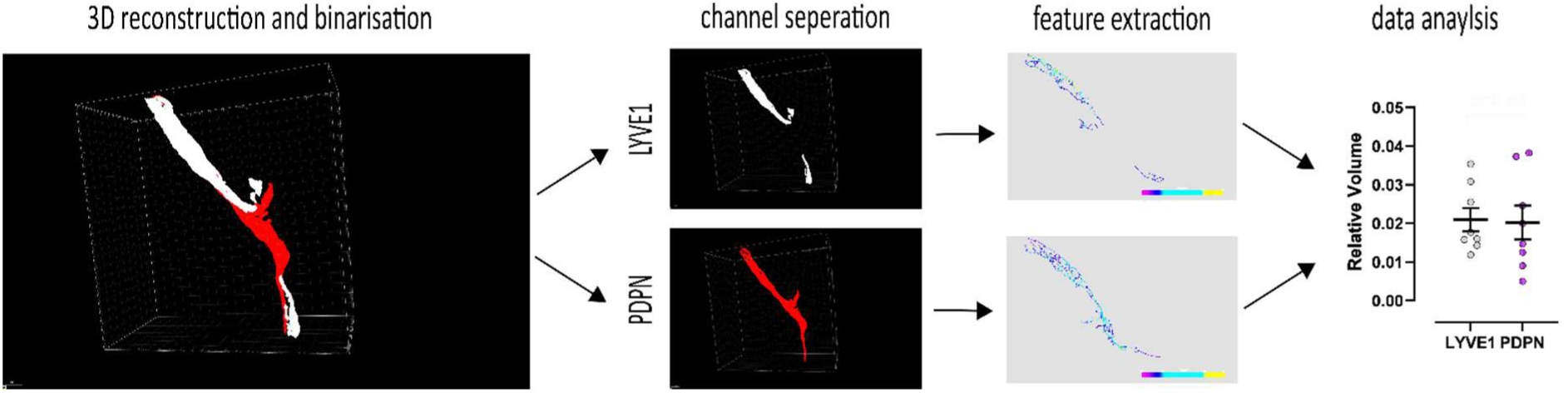
Overview of quantification process. Z-stack data was 3D reconstructed using Imaris software. The Imaris feature “Surface” enabled binarization of desired vessel structures. Thresholds were manually chosen and unwanted structures such as glomeruli in the PDPN channel were actively deleted within the surface feature. This was performed for both channels individually. Following the creation of two separate surfaces for the PDPN and LYVE1 channel, both channels were masked and separately saved as TIFF. Next, vessel volume was extracted using the open-source software VesselVio (Bumgarner and Nelson 2022). Lastly, values for separate channels of one sample were compared, visualised and statistical testing was carried out.

**Supplementary Figure 5:**
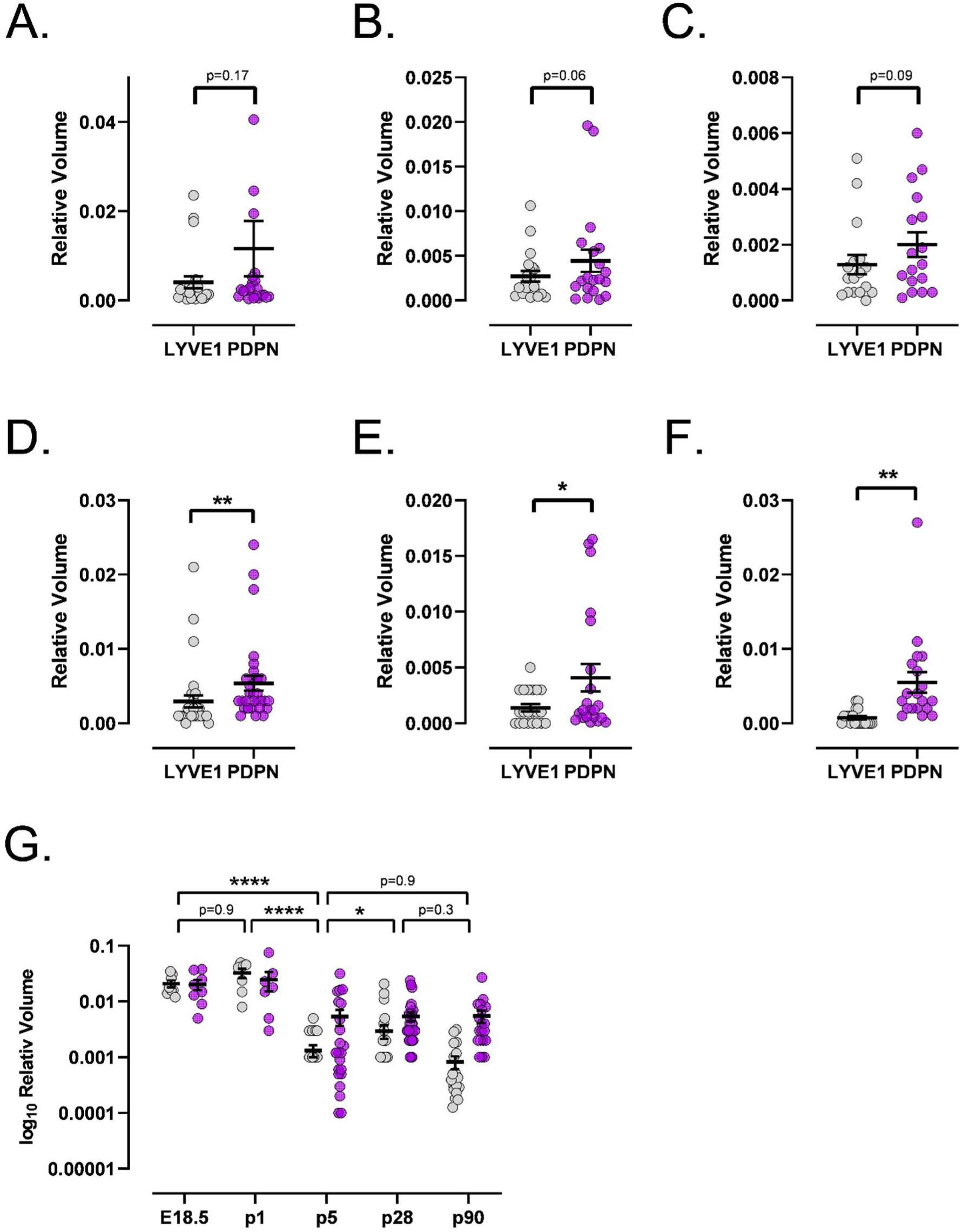
Quantitative analysis of LVYE1^+^ and PDPN^+^ volume per region of interest. Quantitative analysis of the relative volume of LYVE1+ and PDPN+ lymphatic vessels in P28 organs is shown, with each data point representing one region of interest and error bars indicating the standard error of the mean. Vessel volumes were adjusted to the overall sample volume. In P28 skin (n=23), paired Student’s t-test revealed no significant difference between the relative volumes of PDPN and LYVE1 (p=0.169; mean difference = 0.0075, t=1.1421, df=22) (**A**). In P28 heart (n=20), no significant difference was found (p=0.063; mean difference = 0.0017, t=1.976, df=19) (**B**). In P28 lung (n=18), no significant difference was observed (p=0.097; mean difference = 0.0007, t=1.765, df=16) (**C**). In P28 kidney (n=31), a highly significant difference was found (p=0.009; mean difference = 0.0025, t=2.753, df=30). (**D**) In P5 kidney (n=21), a significant difference was observed (p=0.014; mean difference = 0.0027, t=2.684, df=20) (**E**). Note that E18.5 and P1 kidneys were imaged as a whole, so no separate region of interest data is available (see Fig. 3B-C). In P90 kidney (n=19), a highly significant difference was found (p=0.004; mean difference = 0.0047, t=3.321, df=18) (**F**). (**G**) Statistical analysis of LYVE1 volume dynamics throughout development. Nonparametric multiple comparisons for relative contrast effect testing (estimation method = global pseudo ranks, type of contrast = Tukey, confidence level = 95 %) (Konietschke et al. 2015) for five developmental timepoints did find a significant alteration between the timepoints (overall p value=1.02e^−08^, quantile=2.75). As expected, a significant alteration between E18.5 and P1 could not be found (p=0.96, estimator=0.603, lower=0.292, upper=0.848, statistic=0.879). Further, no statistically significant difference could be detected between timepoints P28 and P90 (p=0.34, estimator=0.375, lower=0.225, upper=0.554, statistic=-1.926). Statistical testing did find a significant change between developmental stage P1 and P5 (p=3.48e^−07^, estimator=0.054, lower=0.013, upper=0.196, statistic=-5.006) and P5 and P28 (p=0.038, estimator=0.321, lower=0.186, upper=0.493, statistic=-2.850), confirming a decline in LYVE1^+^ vessels throughout maturation. Further, non-chronological timepoints show highly significant differences, such as E18.5 and P28 (p=1.02e^−08^, estimator=0.053, lower=0.015, upper=0.174, statistic=-5.998), P1 and P28 (p=1.49e^−06^, estimator=0.065, lower=0.016, upper=0.223, statistic=-5.148) or P1 and P90 (p=5.82e^−05^, estimator=0.049, lower=0.010, upper=0.208, statistic=-5.006). No statistically significant change was detected between P5 and P90 (p=0.9, estimator=0.529, lower=0.354, upper=0.697, statistic=0.450).

